# Can pornography be addictive? An fMRI study of men seeking treatment for problematic pornography use

**DOI:** 10.1101/057083

**Authors:** Mateusz Gola, Małgorzata Wordecha, Guillaume Sescousse, Michał Lew-Starowicz, Bartosz Kossowski, Marek Wypych, Scott Makeig, Marc N. Potenza, Artur Marchewka

## Abstract

Pornography consumption is highly prevalent, particularly among young adult males. For some individuals, problematic pornography use (PPU) is a reason for seeking treatment. Despite the pervasiveness of pornography, PPU appears under-investigated, including with respect to the underlying neural mechanisms.

Using functional magnetic resonance imaging (fMRI), we examined ventral striatal responses to erotic and monetary stimuli, disentangling cue-related ‘wanting’ from reward-related ‘liking’ among 28 heterosexual males seeking treatment for PPU and 24 heterosexual males without PPU. Subjects engaged in an incentive delay task in the scanner, in which they received erotic or monetary rewards preceded by predictive cues. BOLD responses to erotic and monetary cues were analyzed and examined with respect to self-reported data on sexual activity collected over the 2 preceding months.

Men with and without PPU differed in their striatal responses to cues predicting erotic pictures, but not in their responses to erotic pictures. PPU subjects when compared to control subjects showed increased activation of ventral striatum specifically for cues predicting erotic pictures but not for cues predicting monetary gains. Relative sensitivity to cues predicting erotic pictures versus monetary gains was significantly related to the increased behavioral motivation to view erotic images (suggestive of higher ‘wanting’), severity of PPU, amount of pornography use per week and number of weekly masturbations.

Our findings suggest that, similar to what is observed in substance and gambling addictions, the neural and behavioral mechanisms associated with the anticipatory processing of cues specifically predicting erotic rewards relate importantly to clinically relevant features of PPU. These findings suggest that PPU may represent a behavioral addiction and that interventions helpful in targeting behavioral and substance addictions warrant consideration for adaptation and use in helping men with PPU.

## Introduction

Pornography consumption has become highly prevalent, in part given Internet availability (Luscombe, 2016). Approximately 70% of males and 20% of females aged 18-30 years use pornography weekly (Hald, 2006). Among teenagers less than 18 years of age, 90% of boys and 60% of girls have used Internet pornography (Sabina *et al*, 2008), with 12% of children having onset of regular consumption below age 12 (Opinium Research, 2014). For most people, pornography viewing is a form of entertainment, but for some individuals problematic pornography use (PPU) accompanied by excessive masturbation promotes seeking of treatment (Gola *et al*, 2016a). Such observations raise multiple scientifically and clinically important questions, including with respect to brain mechanisms related to PPU and their relationships to clinically relevant measures. Given the negative health measures associated with compulsive sexual behavior (CSB) broadly (e.g., childhood sexual trauma and post-traumatic stress disorder (Smith *et al*, 2014)), more research is needed in order to better understand specific forms of CSB like PPU and develop improved intervention strategies (Kafka, 2014; Kor *et al*, 2013; Kraus *et al*, 2016).

The existence and clinical utility of non-substance or behavioral addictions has been debated, with gambling disorder currently being the sole non-substance disorder classified together with substance-use disorders in DSM-5 (American Psychiatric Association, 2013; Holden, 2001). Although a field trial for hypersexual disorder (Kafka, 2010) was conducted, neither this condition or related behaviors such as PPU were included in DSM-5, in part given the relative paucity of data on these behaviors or conditions (Kafka, 2014; Krueger, 2016; Reid *et al*, 2012). Whether excessive and problematic patterns of sexual behavior are best conceptualized within obsessive-compulsive-disorder (OCD), impulse-control-disorder (ICD), behavioral-addiction or other frameworks has been debated (Kafka, 2014; Kor *et al*, 2013; Kraus *et al*, 2016). A recent case series reported that low dose (20 mg/day) of paroxetine treatment (found to be successful in treating OCD (Stein *et al*, 2007)) led to reductions in anxiety and severity of PPU (Gola and Potenza, 2016). Additionally, naltrexone treatment (found to be successful in alcohol-use (Maisel *et al*, 2013) and gambling disorders (Yip and Potenza, 2014) may be helpful for individuals with PPU (Bostwick and Bucci, 2008; Kraus *et al*, 2015). As naltrexone has been proposed to reduce craving through modulating activity in mesolimbic structures (Thompson *et al*, 2000), the ventral striatum may contribute importantly to compulsive sexual behaviors including PPU. Recent MRI studies of men support this hypothesis. Among non-problematic pornography users, an inverse relationship between right caudate volume and frequency of pornography consumption was observed (Kühn and Gallinat, 2014). Increased blood-oxygen-level-dependent (BOLD) responses in the ventral striatum were observed in response to preferred sexual pictures when compared to non-preferred ones, and this activity positively correlated with scores on the Internet Addiction Test Modified for Cybersex (Brand *et al*, 2016). Men with CSB (meeting criteria for hypersexual disorder (Kafka, 2010)) as compared to those without (comparison subjects; CSubs) demonstrated increased striatal reactivity for sexually explicit videos (Voon *et al*, 2014) and decreased functional connectivity between the ventral striatum and prefrontal cortex (Klucken *et al*, 2016). These findings suggest similarities between CSB and addictions.

A prominent model of addiction, the incentive salience theory (IST; (Berridge, 2012; Robinson and Berridge, 1993; Robinson *et al*, 2015), posits that wanting becomes dissociated from liking. The latter is hypothesized to be linked to the *experienced* value of the reward and the former to its *anticipated* value (Robinson *et al*, 2015). ‘Wanting’ is typically evoked by predictive cues associated with reward through Pavlovian learning (Berridge, 2012). Learned cues (conditional stimuli) related to addiction acquire incentive salience, reflected in increased BOLD response in the ventral striatum and increased motivated behavior (i.e. shorter reaction times; RTs) (Berridge, 2012). According to the IST, and consistent with observations in substance addictions and gambling disorder (Robinson *et al*, 2015; Sescousse *et al*, 2013), increased anticipatory ‘wanting’ is dissociated from experienced ‘liking’ in addiction. If PPU share mechanisms with addictions, we anticipate seeing increased BOLD response in the ventral striatum specifically for cues signaling erotic pictures followed by higher motivation to obtain them (measured as shorter RTs) in individuals with PPU compared to CSubs. Increased ‘wanting’ should be unrelated to measures of ‘liking’ in PPU subjects, but not in CSubs.

The current study sought to extend prior studies by examining the neural correlates of sexual and non-sexual images in men seeking treatment for PPU and men without PPU. We further sought to relate the brain activations to clinically relevant features of PPU. No prior neuroimaging studies have examined individuals seeking treatment for PPU. Additionally, as it is important to investigate possible common neural mechanisms of addictions, we investigated cue-induced “*wanting*” of “addiction-related” reward dissociated from “*liking*” aspects. Most studies using visual sexual stimuli do not allow for determination of whether stimuli may represent cues or rewards (Gola, 2016; Gola *et al*, 2016c) and very rarely permit comparisons to other incentives, making it difficult to interpret results with respect to the IST (Berridge, 2012; Gola *et al*, 2016d; Robinson and Berridge, 1993; Robinson *et al*, 2015).

To investigate, we used an incentive delay task (Figure 1) previously used in studies of gambling disorder (Sescousse *et al*, 2013). This task has three important properties; it: 1) disentangles cue- and reward-related phases related to anticipation and outcome, respectively; 2) allows measurement of neural and behavioral indicators of ‘wanting’ (in a cue phase) and ‘liking’ (in a reward phase); and, 3) provides a possibility to compare “addiction-related” stimuli (in this case erotic pictures) with another potent reward (monetary gains). As individuals with gambling disorder expressed higher ventral striatal responses to monetary as compared to erotic cues in the cue phase (Sescousse *et al*, 2013), we hypothesized that men with PPU as compared to those without would demonstrate increased ventral striatal responses for erotic but not monetary cues. We further hypothesized that the degree of ventral striatal activation to erotic cues in the men with PPU would correlate positively with severity of PPU, amount of pornography consumed and frequency of masturbation. Finally according to the IST, we hypothesized that ‘wanting’ in the erotic cue phase would be associated with ‘liking’ in the reward phase in the CSub group but not the PPU group, representing a dissociation between ‘wanting’ and ‘liking’ in PPU.

**Figure 1.**
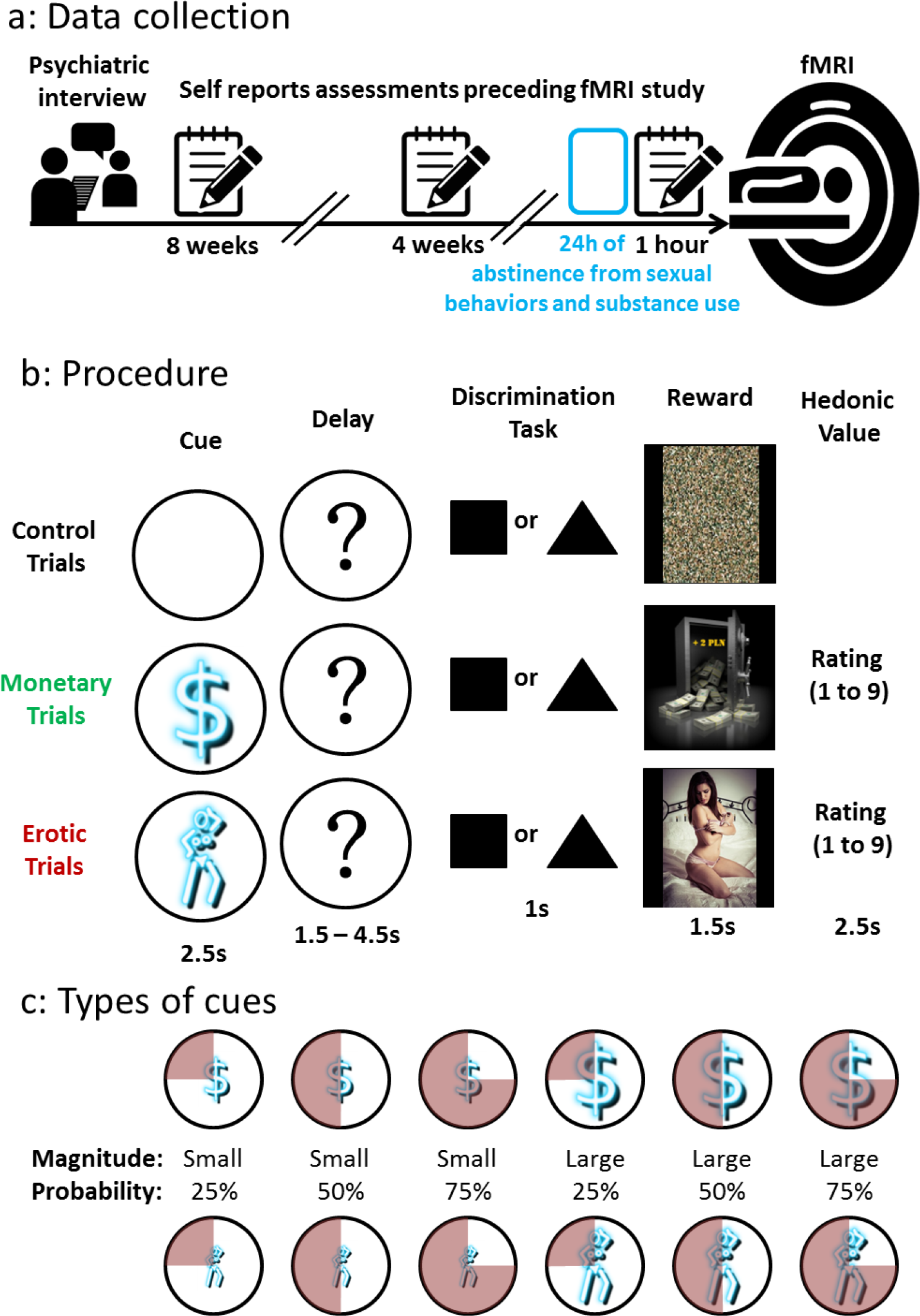
Data collection and experimental procedure. a: After psychiatric assessment, subjects who met study criteria (see Methods) completed questionnaires assessing self-reported sexual behavior and substance use in the weeks preceding fMRI. **b: Incentive delay task** used in the fMRI session. Subjects first saw a cue informing them about the type (pictogram with a $ or a woman), magnitude (size of pictogram) and probability (pie chart) of an upcoming reward. Examples of all possible cases of cues are presented in fig. 2c. An empty circle was used to signal control trials with a 100% likelihood of getting no reward (fig. 1b, top). Next, the cue was replaced by a question mark, symbolizing a delay period during which a pseudorandom draw was performed according to the previously displayed probability. Following this anticipation phase, participants had to perform a target discrimination task within 1 s. The target was either a triangle (left button press required) or a square (right button press required). If they answered correctly within less than 1 s, they were then allowed to view the outcome of the pseudorandom draw. Reaction times were later used as an index of motivation. In rewarded monetary trials (following the cue with pictogram of a $; fig. 2b, middle) subjects saw a monetary amount displayed on a safe. In rewarded erotic trials (following cue with the pictogram of woman; fig. 2b, bottom) subjects saw an erotic picture. After each reward outcome, subjects had to provide a hedonic rating on a continuous scale (1-don’t like it to 9-like it very much). In non-rewarded and control trials, subjects saw a scrambled picture (bottom). **c: Types of cues**. Both monetary and erotic cues provided information about the magnitude and probability of reward. For erotic rewards, a small magnitude was always predictive of pictures of women in lingerie or swimming suits, and a large magnitude was always predictive of explicit pictures of woman in postures inviting sexual activity. For monetary cues, a small magnitude was predictive of gains ranging from 1 to 3 PLN (proximately .25-.75 EUR), while a large magnitude was predictive of gains ranging from 6 to 8 PLN (1.5 – 2 EUR). The probability of obtaining a reward after a correct and fast (<1 s) response was indicatied by a background pie chart representing probabilities of 25%, 50% or 75%. All monetary gains were paid to the participants at the end of the experiment. Credits for the sample photo: Lies Thru a Lens, CC BY 2.0. For license terms see: CC BY 2.0.

**Table 1.** Subject characteristics

|  | Control Sub (N=24)<br>Mean (SD) | PPU (N=28)<br>Mean (SD) | p value |
| --- | --- | --- | --- |
| Age | 30.49 y.o. (7.55) | 30.96 y.o. (6.51) | n.s.<sup>A</sup> |
| Monthly income<sup>B</sup> | 3360 PLN (2264)<sup>C</sup> | 3463 PLN (2491)<sup>C</sup> | n.s.<sup>A</sup> |
| Average pornography consumption per week<sup>D</sup> | 50.77 min (42.6) | 287.87 min (258.4) | <.001 |
| Longest time of pornography consumption in one day<sup>E</sup> | 70.55 min (52.8) | 284.74 min (321.3) | =.002 |
| Average frequency of masturbation per week<sup>D</sup> | 2.37 (1.46) | 5.66 (3.04) | <.001 |
| Max number of masturbations in one day | 3.10 (1.17) | 5.21 (2.75) | <.001 |
| Sex Addiction Screening Test –Revised (SAST-R) | 2.67 (2.12) | 11.46 (4.95) | <.001 |
| Brief Pornography Screener (BPS)<sup>F</sup> | 2.88 (3.24) | 5.91 (2.96) | =.005 |
| Sexual Arousalability Inventory (SAI) | 83.39 (19.27) | 88.64 (19.89) | n.s.<sup>A</sup> |
| Obsessive Compulsive Inventory –Revised (OCI-R)<sup>F</sup> | 14.13 (6.73) | 15.77 (10.10) | n.s.<sup>A</sup> |
| UPPS-P Impulsive Behaviour Scale (UPPS-P)<sup>F</sup> | 130.73 (24.97) | 131.87 (25.37) | n.s.<sup>A</sup> |
| Eysenk's Impulsivity Inventory – Impulsivity (IVE-I) | 5.95 (3.85) | 6.95 (5.01) | n.s.<sup>A</sup> |
| Eysenk's Impulsivity Inventory – Risk Taking (IVE-R) | 9.41 (4.01) | 9.9 (3.96) | n.s.<sup>A</sup> |
| State-Trait Anxiety Inventory – State (STAI-S) | 33.79 (6.35) | 37.69 (9.36) | n.s.<sup>A</sup> |
| State-Trait Anxiety Inventory – Trait (STAI-T) | 37.83 (6.96) | 43.60 (9.88) | =.022 |
| Substance use prior to fMRI session | Number of subjects using substances within the last:<br>24 hours / 2-7 days / 8-90 days / > 90 days / never |  |  |
| Tobacco | 3 / 2 / 4 / 10 / 5 | 3 / 6 / 5 / 11 / 6 |  |
| Alcohol | 0 / 11 / 11 / 2 / 0 | 0 / 11 / 11 / 6 / 0 |  |
| Marijuana | 0 / 1 / 3 / 12 / 7 | 0 / 3 / 5 / 14 / 6 |  |

<sup>A</sup> Non-significant: $p > .2$
<sup>B</sup> Income after taxes
<sup>C</sup> 1 PLN is approximately .25 EUR. Average monthly income after taxes in Poland in 2014 was 2865 PLN or about 716 EUR.
<sup>D</sup> Averaged across three self-reports about 8<sup>th</sup>, 4<sup>th</sup> and last week before the fMRI recording (see Figure 1a) as all assessments at the 3 timepoints were highly related to each other within each domain (pornography use measure 1 and 3: $R = .871$ ; $p < .01$ , and masturbation: $R = .792$ ; $p < .01$ )
<sup>E</sup> Longest time spent on pornography consumption within 24 hour period (one day)
<sup>F</sup> These questionnaires were administered during the 2<sup>nd</sup> assessment (4 weeks before fMRI session). Due to the time needed for development and validation of Polish language versions, these questionnaires were added during the study and 16 CSubs and 22 PPU subjects completed them. All other questionnaires were
administered during the 1<sup>st</sup> assessment in all subjects (8 weeks before fMRI session). Sexual activity was assessed during all 3 assessments.

References for questionnaires: SAST-R (Carnes *et al*, 2010; Gola *et al*, 2016b); BPS (Kraus et al., under review; see Supplementary Materials); SAI (Gola *et al*, 2015; Hoon *et al*, 1976); OCI-R (Foa *et al*, 2002); UPPS-P (Poprawa, 2016; Whiteside and Lynam, 2003); IVE-I/R (Jaworowska, 2011); STAI-S/T (Sosnowski and Wrześniewski, 1983; Spielberger, 2010).

## Methods

### Participants

Fifty-seven heterosexual males (age range 18-48 years) participated in the fMRI study. These included thirty-one men seeking treatment for PPU (meeting criteria of hypersexual disorder (Kafka, 2010)) and without other psychiatric diagnoses and twenty-six CSubs with comparable ages and incomes, also without psychopathology. All subjects were medication-free. Three PPU subjects and two CSubs were excluded from analysis due to extensive head movement (more than 7mm). Characteristics of the remaining 28 PPU subjects and 24 CSubs are presented in Table 1. All participants were financially compensated based on their winnings accumulated during the experimental procedure (M=184.84 PLN; SD=21.66; approximately 46 EUR). Details of recruitment and anonymity procedures are presented in Supplementary materials. All research procedures were approved by the Ethical Committee of Institute of Psychology, Polish Academy of Science. All subjects provided written informed consent.

### Recruitment

Subjects were recruited among men seeking treatment for PPU in two clinics in Warsaw. After the initial interview, patients were screened for sexual orientation (Inclusion criteria: exclusively or predominantly heterosexual on the Kinsey Scale (Kinsey *et al*, 1948; Wierzba *et al*, 2015) and no history of alcohol abuse (scores <7 on the Alcohol Use Disorder Identification Test (Saunders *et al*, 1993) or gambling problems (scores <3 on the South Oaks Gambling Screen (Lesieur and Blume, 1987). Patients meeting these criteria and having no contraindication for MRI were screened with the SCID-I (First and Gibbon, 2004) for OCD, ICDs, mood disorders, mania, anxiety disorders, psychotic disorders, history of substance abuse/dependence and hypersexual disorder according to criteria proposed by Kafka (Kafka, 2010). Only men meeting criteria for hypersexual disorder and none of the other above mentioned disorders were invited to participate in a 2-month self-assessment study involving completing web-based questionnaires (approximately 8 weeks, 4 weeks, and one hour before the fMRI session; Figure 1) and fMRI.

CSubs were recruited through web-based announcements advertising the study as a survey on Internet pornography use (to avoid primary monetary motivations). Among 213 men, we selected 26 heterosexual individuals matched by age (the same year of birth), income (+/- 15%) and handedness to each PPU subject. All CSubs had used pornography at least once in the preceding year, but had never experienced it as a problematic behavior. All CSubs followed the same procedures as PPU subjects.

### Anonymity

To ensure the anonymity of PPU individuals, we applied a double-blind approach in that the research team in the laboratory had no access to the data gathered by the recruitment and assessment team and thus did not know subjects’ group (PPU, CSub) identities. Each subject received an alphanumeric code to maintain anonymity at the data analysis level. We informed subjects about these procedures.

### Questionnaire assessments

In self-assessments preceding fMRI (Figure 1a), subjects were asked to report their sexual activity during the week (see Table 1). During this phase, we also collected questionnaire measurements for independent verification of screening accuracy and assessment of additional data (as presented in Table 1 and described in detail in the Supplementary Materials).

### Incentive delay task

We used the same procedure described in detail in previous studies (Sescousse *et al*, 2010, 2013), schematized in Figure 1b, and described in Supplementary Materials, with modifications related to the amount of monetary gains. In the original studies, subjects were informed that they would receive a sum of rewards from one randomly chosen experimental block (out of four) (Sescousse *et al*, 2013). In our study, subjects were told they would receive the exact sum of all monetary gains (M=184.84 PLN, which was approximately 5.5% of monthly salary after taxes). The task permits modeling of events theoretically related to ‘wanting’ (cues) and ‘liking’ (rewards).

### MRI data acquisition

MRI data acquisition was conducted at the Laboratory of Brain Imaging, Neurobiology Center, Nencki Institute of Experimental Biology on a 3-Tesla MR scanner (Siemens Magnetom Trio TIM, Erlangen, Germany) equipped with a 12-channel phased-array head coil. Functional data were acquired using a T2*-weighted gradient echo-planar-imaging (EPI) sequence with the following parameters: repetition time=2500ms, echo time=28ms, flip angle=90°, in plane resolution=64×64 mm, field of view=224 mm, and 35 axial slices with 3.5 mm slice thickness with no gap between slices. Each of the four functional runs consisted of 286 volumes. Field mapping was done based on prior methodology (Jezzard and Balaban, 1995) using double-echo FLASH (echo time 1=4.92 ms, echo time 2=7.38 ms time repetition=600 ms) with the same spatial properties as the functional scans. Detailed anatomical data were acquired with a T1-weighted sequence (repetition time = 2530 ms, echo time=3.32 ms). Head movements were minimized with cushions placed around the participants’ heads. Subjects were asked to refrain from any psychoactive substance use and sexual activity during the 24 hours preceding fMRI.

### fMRI Analysis

In line with previous studies (Sescousse *et al*, 2010, 2013), in the first-level analysis we modelled brain responses during the cue-anticipatory phase and the reward-outcome phase. We modelled separately 13 cue conditions: low/high erotic cue and low/high monetary x 3 probability categories – 25%, 50% and 75% (giving 6 erotic-cue categories and 6 monetary-cue categories, as presented in Figure 2c), and one control cue. Brain responses during the reward-outcome phase were modelled as events time-locked to participants’ responses in the discrimination task (Figure 1b). Rewards were displayed only after correct responses with 25%, 50% or 75% probabilities (Figure 1c), which gave 5 conditions (erotic reward/lack of erotic reward, monetary reward/lack of monetary reward, lack of reward following control cue) without hedonic ratings. Details of signal preprocessing are provided in Supplementary Material.

**Figure 2.**
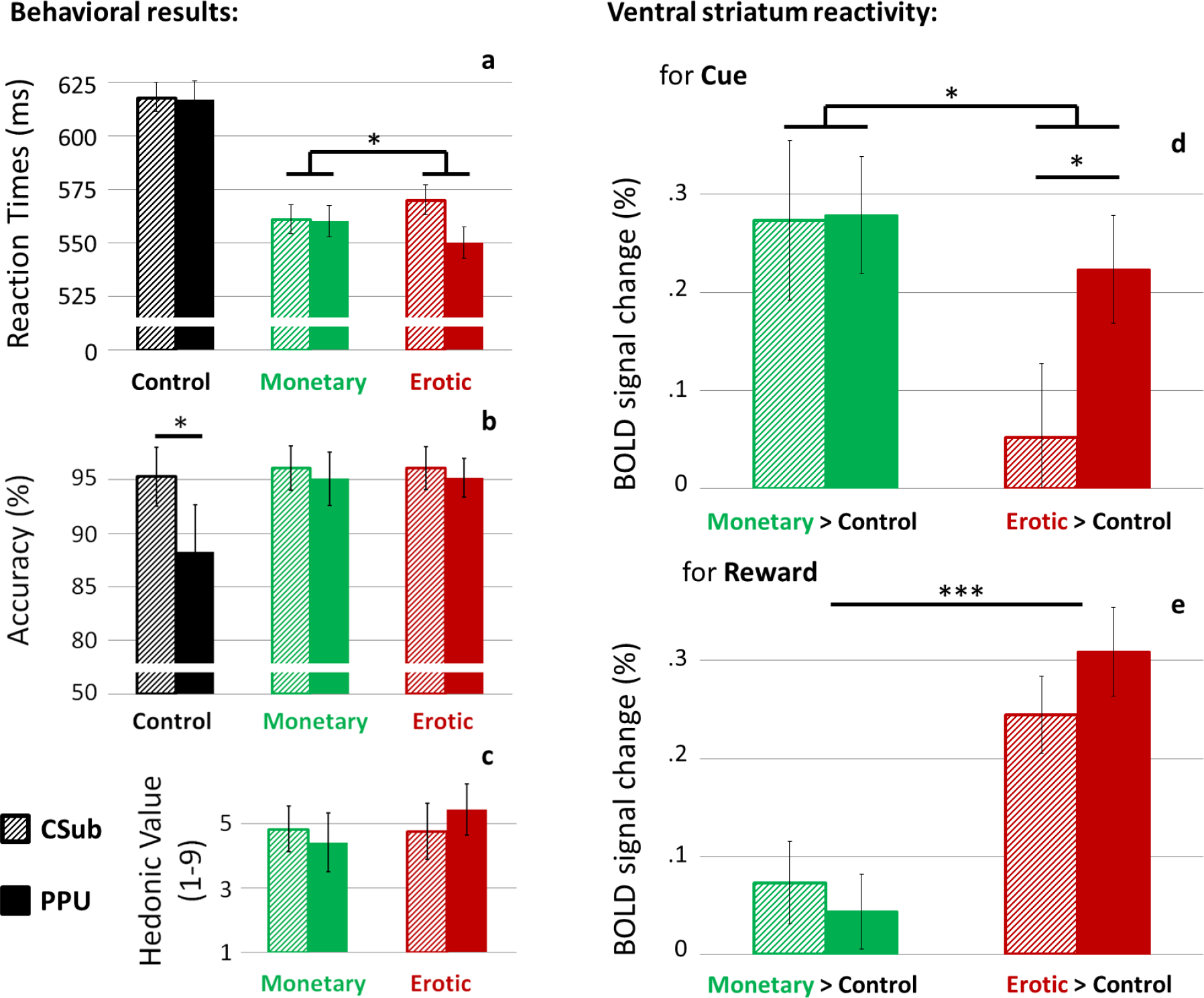
Behavioral and neuroimaging results. **A:** Comparison of RTs in the discrimination task (see Figure 1b). **B:** Comparison of accuracy. **C:** Comparison of hedonic value ratings (see Figure 1b). **D:** Comparison of BOLD signal change in the ventral striatum for cue presentation (BOLD signal averaged across 2 *a priori* defined regions of interests in the left and right brain hemisphere: 8mm spheres centered around: Left: x=-12, y=10 z=-6 Right: x=12, y=10, z=-4). **E:** Comparison of BOLD signal response in the ventral striatum for reward presentation. All post-hoc tests were done with Bonferroni correction for multiple comparisons. Error bars indicates SEM. *p<.05; **p<.01; ***p<.001.

### Regions of interests

We used a striatal ROI defined *a priori* based on a previous meta-analysis of reward anticipation (Liu *et al*, 2011); 8mm spheres were centered around: Left: x=-12, y=10 z=-6, Right: x=12, y=10, z=-4). To focus on our hypothesis and keep this manuscript concise, we present only analyses using the *a priori* defined ROI of the ventral striatum. For control purposes, we also defined an ROI in Heschl’s gyrus, where according to our predictions, no group or condition differences were observed. The ROI was based on the corresponding mask in the AAL atlas taken from the WFU PickAtlas toolbox (version 3.0.5). The percent signal change was calculated with the MarsBaR toolbox (http://marsbar.sourceforge.net). We report significant brain activations within the ROI (Figure 2, 3, 4, S1) that survived family-wise-error (FWE) correction for multiple comparisons using small-volume correction (*P*_*SVC-FWE*_ < 0.05). Due to the very similar effects for left and right ROIs, we present only results averaged across hemispheres.

### Statistical analysis

For statistical analyses, IBM SPSS 22 (IBM Corp. Released 2013, Armonk, NY: IBM Corp) and MATLAB R2014a (The Math-Works Inc., Natick, MA, USA) were used. Due to high correlations across time (at the three time points: 8 weeks, 4 weeks and 1 day before the fMRI) within self-assessed pornography use (R=.871; p<.01) and masturbation (R=.792; p<.01), we computed average scores across time for each variable (Figure 1a; Table 1). For testing group-by-trial-type interactions and main effects of group and trial type in BOLD signal from the ROIs, General Linear Models (GLMs) and Fisher’s F tests were used (ANOVA with trial type, magnitude of cue, probability of cue as within-subject factors and group as a between-subject factor; Figures 2, 4, S1). The same GLMs were used for analysis of reaction times (RTs; Figures 2, 3, S1, S2), accuracy and hedonic value (Figures 2, S2). All *post hoc* comparisons were conducted with Bonferroni-Holms correction. Correlations between BOLD signal and measures of symptoms were computed only for measures significantly differentiating PPU subjects from CSubs (SAST-R, BPS, amount of pornography use and masturbation). Due to the discrete thresholding of SAST-R scores and skewedness of distributions of the three other measures, Spearman’s Rho was used to compute covariance.

## Results

### Between-group differences

Men seeking treatment for PPU and CSubs did not differ in age, income, impulsivity or compulsivity (Table 1). Although no subjects met criteria for anxiety disorders, men with PPU as compared with CSubs exhibited higher trait anxiety (t(50)=2.37; p=.022; STAI-T). No difference in self-declared sexual arousability (SAI) was observed; however, men with PPU as compared with CSubs showed higher sexual arousability for compensatory activities such as pornography use and masturbation on the corresponding SAI subscale (t(46)=3.348; p=.002).

As anticipated, men with PPU as compared with CSubs demonstrated higher scores on the Sex Addiction Screening Test–Revised (SAST-R; t(50)=8.539; p<.001) and Brief Pornography Screening Test (BPS; t(36)=2.998; p=.005) and reported more pornography use (t(50)=3.776; p<.001) and more frequent masturbation (t(50)=5.042; p<.001) during the weeks preceding fMRI.

Both groups had similar accuracy for monetary and erotic trials (Figure 2b) and obtained similar amounts of monetary wins (CSubs: M=187.41 PLN; SD=22.83, PPU subjects: M=182.14 PLN; SD=20.56).

### Behavioral results

We analyzed RTs, accuracy and hedonic values ratings. A significant group-by-cue type interaction on RTs was observed (F(1,50)=5.112; p=.028). The shortest RTs were observed in men with PPU during erotic trials (Fig. 2a). Main effects of group (F(1,50)=.223; p=.639) or cue type (F(1,50)=.390; p=.535) were insignificant. We observed a significant main effect of cue magnitude on RTs (F(1,50)=42.152; p<.000001; Figure 3a). RTs in trials with cues predicting large rewards were shorter. We found a significant interaction between the type and magnitude of cue (F(1,50)=7.416; p=.009). As there were no interactions of probability and group, both in RTs (F(2,50)=1.132; p=.331) and BOLD response (F(2,50)=2.046; p=.135), we present these results in Supplementary Material (Figure S1). There was also a main effect of cue probability on RTs (F(2,50)=20.671; p<.000001), but no group-by-probability interaction (F(2,50)=1.132; p=.331); thus, we present these results in Supplementary Material (Figure S1).

**Figure 3.**
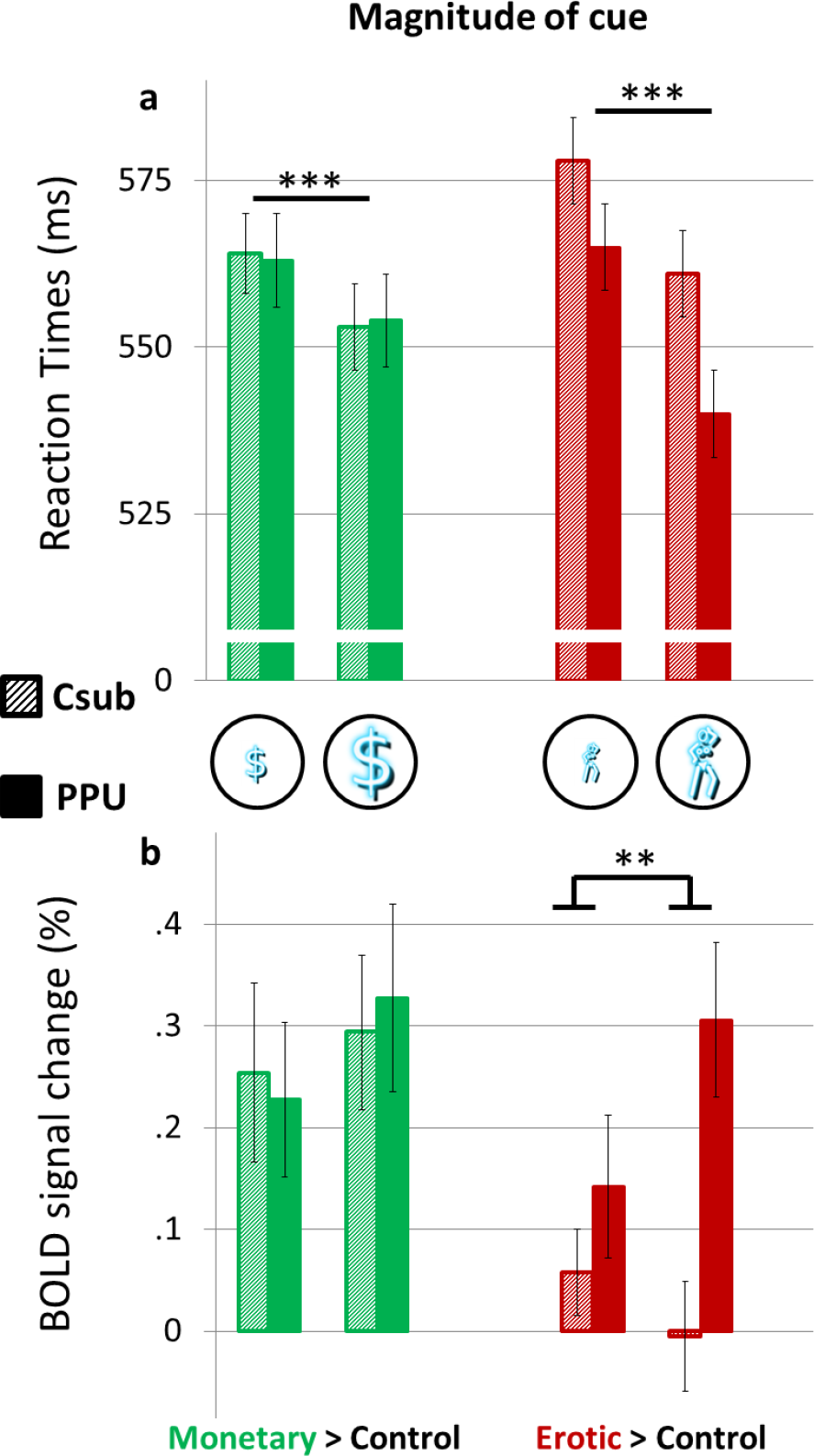
Between-group differences in ventral striatal reactivity for cues predicting small and large magnitudes of monetary (green) and erotic (red) rewards. **A:** Comparison of RTs for cues predictive of small and large rewards. **B:** Comparison of BOLD signal response in the ventral striatum (the same ROIs as in fig. 2) for presentations of cues with different magnitudes. Error bars indicates SEM. **p<.01; ***p<.001.

Analysis of accuracy revealed interesting results. Although there was no group (F(1,50)=.619; p=.435) or group-by-cue-type interaction (F(1,50)=.002; p=.969) in accuracy for reward-related cues (monetary and erotic), there was a trend toward an effect of group-by-cue-type interaction taking into account non-rewarded control cues (F(2,100)=3.014; p=.054). This pattern appears driven by decreased accuracy of PPU men (compared to CSubs) on control (non-rewarded) trials (t(50)=2.084; p=.045). This result shows that while PPU and CSubs have comparable accuracy in trials providing chance for a reward (monetary and erotic), PPU men demonstrate decreased accuracy in non-rewarded (control) trials (Fig. 2b)

No differences were observed in hedonic value ratings (F(1,50)=.187; p=.667; Fig. 2c), which may be considered declarative measure of liking.

### Neuroimaging results

Like with RTs, cue-related reactivity of the ventral striatum demonstrated a group-by-cue-type interaction (F(1,50)=6.886; p=.011; Fig 2d). Men with PPU and CSubs differed significantly in reactivity for erotic (t(50)=2.624; p=.011) but not monetary (t(50)=.047; p=.963) cues. During the reward-processing phase, a main effect of cue type was observed (F(1,50)=44.308; p<.001), but no between-group differences were observed (F(1,50)=.061; p=.806). Also no group-by-reward-type interaction (F(1,50)=.2.054; p=.158) was observed. These results indicate that PPU men and CSubs differ during the cue phase, but demonstrate similar ventral striatal reactivity in the reward phase, with results showing a significant three-way group-by-cue-type-by-processing-phase (cue vs reward) interaction (F(1,50)=5.438; p=.024). The results are consistent with a dissociation between brain measures of ‘wanting’ and ‘liking’ of erotic stimuli in PPU subjects, but not in CSubs.

We also observed a significant group-by-cue-type-by-magnitude interaction in ventral striatal BOLD signal change during the cue phase (F(1,50)=5.432; p=.024). While for monetary cues there was no group-by-magnitude interaction (F(1,50)=.613; p=.482) or main effect of group (F(1,50)=.002; p=.963), the group-by-magnitude interaction (F(1,50)=.8.273; p=.007) and main effect of group (F(1,50)=5.914; p=.019) were significant for erotic cues. The findings indicate that stronger BOLD response in the ventral striatum for erotic cues among PPU men (compared with CSubs) is modulated by magnitude of erotic cue (Figure 3b). The whole-brain BOLD response for the second-level contrast _PPU_(high>low magnitude) > CSubs(high>low magnitude) is presented in Figure S3. We also observed a main effect of cue probability on ventral striatal BOLD signal change (F(2,50)=5.379; p=.006). As there were no group-by-probability effects (F(2,50)=2.046; p=.135), we present these results in Supplementary Materials (Figure S1).

### Relationships between behavioral and neuroimaging results and clinical features of PPU

In line with previous work (Sescousse *et al*, 2013, 2015), we computed for each subject the differential reactivity to monetary versus erotic cues by subtracting the corresponding striatal BOLD responses. We also calculated a relative-motivation index measured as the difference in mean RTs for monetary and erotic trials. The brain-behavior correlation between these two measures was strongly significant (R=.76; p<.0001, Fig. 4a). Next, we examined how individual differences in ventral striatal reactivity for erotic versus monetary cues related to four measures differentiating both groups (Table 1): severity of CSB symptoms measured with the SAST-R (Rho=.31; p=.01; Fig. 4b), pornography craving measured with the BPS (Rho=.264; p=.055), amount of pornography consumption (Rho=.305; p=.015; Fig. 4c) and number of masturbations per week (Rho=.296; p=.018; Fig. 4d). Bonferroni-Holms correction for multiple comparisons was used. We also examined how individual differences in ventral striatal reactivity for erotic versus monetary cues related to masturbation (Rho=.313; p=.034) and amount of pornography consumption (Rho=.47; p=.054) only among PPU subjects.

In the next step, we checked if an analogous index of liking would be related to behavioral measures. For this purpose, we calculated an individual relative-liking index measured as differences in ventral striatal reactivity for erotic and monetary rewards. We related this index to individual differences in ratings of rewards’ hedonic values and the same four measures differentiating both groups as above. None of the correlations were significant.

## Discussion

Our results, in line with the IST (Berridge, 2012; Gola *et al*, 2016d; Robinson and Berridge, 1993; Robinson *et al*, 2015), indicate that men seeking treatment for PPU when compared to CSubs show increased ventral striatal reactivity for cues predicting erotic pictures (but not for cues predicting monetary gains). Such increased striatal reactivity for cues predicting erotic content is followed by higher motivation (reflected in shorter RTs) to view erotic rewards (Figure 2d, 2a and 4a). Consistent with the IST (Robinson and Berridge, 1993; Robinson *et al*, 2015), these results suggest increased ‘wanting’ evoked specifically by an initially neutral cue predictive for erotic rewards. Also as predicted by the IST, a neural indicator of ‘wanting’ (BOLD response in the ventral striatum) is dissociated from measures of ‘liking’ among PPU men but not CSubs, and this was reflected in three-way interactions between incentives (monetary versus erotic), group (CSubs versus PPU), and experimental phase (cue versus reward). In other words, subjects who seek treatment for PPU expressed higher motivational behavior for pornography cues predictive of erotic content. This motivated behavior (‘wanting’, probably related to the expectation of highly rewarding value of pornography) is dissociated from actual ‘liking’: PPU subjects did not differ from CSubs in BOLD response for erotic pictures (reward phase) or hedonic values ratings (Figure 2c and 2e). Moreover, the differential striatal reactivity to erotic versus monetary cues (but not rewards) was related not only to indicators of motivated behaviors during the study (RTs; Figure 4a), but also to the severity of CSB (measured with the SAST-R; Figure 4b), amount of pornography consumption and frequency of masturbation (Figure 4c and 4d) reported during the 2 months preceding fMRI.

**Figure 4.**
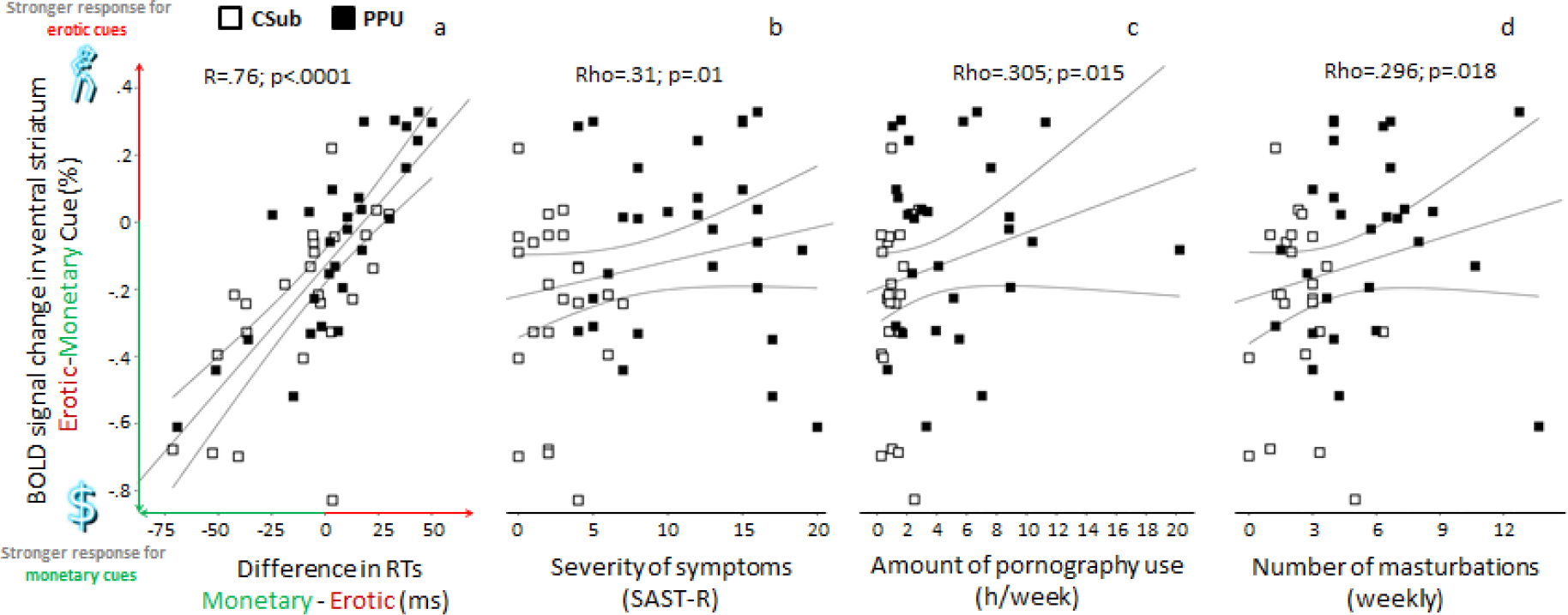
Correlations of ventral striatal cue-reactivity with amount of pornography use, frequency of masturbation and clinical features of PPU. The correlations between differential striatal reactivity to monetary versus erotic cues and **A:** relative-motivation index measured as difference between RTs for monetary - erotic trials; **B:** severity of CSB measured by the Sexual Addictions Screening Test – Revised (20 points scale, which was not used at recruitment phase); **C:** average amount of pornography consumption per week, and **D:** frequency of masturbation per week. Error lines depicts 95% confidence intervals. Bonferroni-Holms correction was used for multiple comparisons.

This pattern of increased cue-related ‘wanting’ dissociated from reward-related ‘liking’ resembles findings in addictions (Robinson *et al*, 2015); (Sescousse *et al*, 2013). Specific cues (predictive for addiction-related rewards) evoke activations of brain-reward systems associated with striatal responses (Flagel *et al*, 2011; Oei *et al*, 2012; Robinson and Berridge, 1993; Smith *et al*, 2011) and motivations to approach rewards, but experienced hedonic value (Berridge, 2012; Robinson *et al*, 2015) or striatal response for reward (Flagel *et al*, 2011) are not proportional to ones evoked by the preceding cue. These findings are consistent with an impaired mechanism of updating cue-related predictions about expected values of erotic stimuli, similar to mechanisms proposed for substance-use disorders (Parvaz *et al*, 2015; Tanabe *et al*, 2013), although this possibility warrants direct investigation. Given the role of the ventral striatum in reward anticipation (Balodis and Potenza, 2015), initially neutral stimuli (akin to cues introduced in our experimental procedure) may become for men with PPU powerful incentives under the circumstances of pairing them with erotic images. Our results show that individuals with PPU are much more sensitive (then CSubs) for cues signaling erotic rewards (Figure 2d) and the magnitude of the expected erotic reward further modulates the ventral striatal reactivity in men with PPU, which does not happen among CSubs (Figure 4). These results are in line with recent studies showing stronger effects of conditioning for cues predicting explicit sexual content among individuals with CSB compared with CSubs (Banca *et al*, 2016; Klucken *et al*, 2016). Along with these and other studies (Gola *et al*, 2016d; Mechelmans *et al*, 2014), our results suggest that conditioned stimuli associated with erotic rewards may overshadow motivational values of alternate sources of reward in men with PPU, eventually leading to PPU. However, longitudinal studies are needed to examine this hypothesis.

Our results show that PPU is related to alterations in motivational processes. Besides increased striatal response for cues predicting erotic rewards and accompanying increases in motivated behaviors (RTs to erotic cues), men with PPU also exhibited a decrease in motivated behaviors for non-rewarded trials (lower accuracy, fig 2b) when compared to CSubs. This finding raises questions whether men with PPU may have more generalized impairments of reward processing, in line with a reward deficiency syndrome theory (Comings and Blum, 2000), which would predict decreased striatal reactivity and accompanying hedonic values for both types of rewards (erotic and monetary) in the reward phase. Here we show that this is not the case; men with and without PPU do not differ in striatal reactivity and hedonic values either for erotic or monetary rewards. The key difference between these two groups is in striatal reactivity (and accompanying behavioral reactions) in response for cues.

In our study, we have made the assumption that RTs (Figure 4a) directly reflect motivation. Such a coupling has been assumed in several previous studies (Clithero *et al*, 2011; Sescousse *et al*, 2015), and is grounded in computational work linking motivation and vigor via dopamine (Niv *et al*, 2007). Also, we think that two aspects of our design help narrowing the interpretation of RTs in terms of motivation rather than response vigor. The first is the motor component of our task was not a simple target detection task but a discrimination task with two possible button responses, precluding the interpretation of differences in RTs in terms of motor preparation. In addition, the use of the monetary cue condition can be regarded as a “baseline” condition controlling for group differences in response vigor or motor activation. Yet, in the absence of self-report “wanting” ratings, we cannot rule out the potential influence of motor activation on RTs. Complementary procedures such as preference tasks could be used in future studies to refine the interpretation of the current results. Additionally, given possible carry-over effects, we cannot entirely exclude the potential influence of rewards on subsequent cue processing, although we believe that our counter-balanced task design helps to mitigate against this possibility.

Despite the above-mentioned limitations, our findings indicate that increased cue-reactivity among men with PPU is not a general dysfunction, but is related to cues predictive of erotic but not monetary rewards. This selective mechanism of increased reactivity for erotic but not monetary cues among men with PPU speaks in favor of the IST (Robinson and Berridge, 1993) rather than more generalize impairments of reward processing proposed in example by theoretical frameworks such as the reward deficiency syndrome (Comings and Blum, 2000). However, the resolution of fMRI may not permit the possible resolution of hedonic hotspots related to wanting or liking signals. Lack of between-group differences in responses to monetary cues and compulsivity (measured with the OCI-R) also speaks against conceptualization within an OCD framework (Kor *et al*, 2013), as OCD patients present decreased striatal BOLD response for cues predictive for monetary gains when compared to CSubs (Figee *et al*, 2011). However, it is important to note that we were excluding all subjects with comorbid psychiatric disorders (approximately 50% of treatment-seeking individuals were excluded for this reason during the initial screening procedure), so our conclusion about a lack of generalized reward processing impairment may not generalize to men with PPU and comorbid disorders. Despite this limitation, the exclusion of individuals with PPU and co-occurring psychiatric disorders permitted for a more focused study on mechanisms underlying PPU and excludes possible effects of psychopathology. Additional limitations include the exclusion of women, and future studies should examine the extent to which the findings extend to women with PPU. Additionally, future studies should examine how neurobiological and clinical measures might relate to treatment outcomes for individuals with PPU.

## Conclusions

Men with PPU showed increased activation of the ventral striatum specifically for cues predicting erotic but not monetary rewards. In PPU subjects, this brain activation was accompanied by measures suggesting increased behavioral motivation to view erotic images (higher ‘wanting’). Ventral striatal reactivity for cues signaling erotic pictures (but not for erotic pictures *per se*) was significantly related to severity of CSB, amount of pornography use per week and frequency of masturbation. The findings suggest similarities between PPU and addictions and an important role for learned cues in PPU. Identifying PPU-related triggers and targeting the dissociation of learned cues from problematic behaviors may be useful in the treatment of PPU. Future studies should examine specific treatments, as well as determine the prevalence and clinical correlates of PPU, and identify predisposing factors for PPU.

## Acknowledgements

We are grateful to all participants who agreed to be involved in this study; M. Skorko and the VR Lab (http://vrlab.edu.pl/) for the possibility of using the GEx platform to collect the questionnaire data; M. Wilk, P. Winkielman, E. Kowalewska, M. Bielecki, P. Holas, Ł. Okruszek, Ł. Głowacki, W. Ciemniewski and D. Baran for help and comments. This study was supported by the Polish National Science Centre, OPUS grant, number 2014/15/B/HS6/03792 (M. Gola). M. Gola was also supported by the Polish Ministry of Science scholarships (1057/MOB/2013/0 and 469/STYP/10/2015) scholarship Start of Foundation for Polish Science and scholarship of The Kosciuszko Fundation. G. Sescousse was supported by a Veni grant from the Netherlands Research Organization (NWO). M. Potenza is supported by the National Center on Addiction and Substance Abuse and the National Center for Responsible Gaming. The views presented in the article do not necessarily reflect those of the funding agencies and rather reflect those of the authors.

## Authors Contributions

M.G., G.S., A.M designed the project. M.G., G.S. and B.K. prepared experimental procedures. M.G. and M.L-S., recruited subjects and collected clinical data. M.Wo. conducted the experiments. M.G., M.Wo., B.K., M.Wy., J.R. and A.M performed the statistical analysis. M.G., G.S., M.P. and M.Wo analyzed the findings. M.G., G.S. and M.P. wrote the manuscript.

